# Time-of-day effects in resting-state fMRI: changes in Effective Connectivity and BOLD signal

**DOI:** 10.1101/2020.08.20.258517

**Authors:** Liucija Vaisvilaite, Vetle Hushagen, Janne Grønli, Karsten Specht

## Abstract

The current project explored the hypothesis that time-of-day dependent metabolic variations may contribute to reduced reliability in resting-state fMRI studies. We have investigated time-of-day effects in the spontaneous fluctuations (>0.1Hz) of the blood oxygenation level dependent (BOLD) signal. Using data from the human connectome project (HCP) release S1200, cross-spectral density dynamic causal modelling (DCM) was used to analyze time-dependent effects on the hemodynamic response and effective connectivity parameters. Hierarchical group-parametric empirical Bayes (PEB) found no support for changes in effective connectivity, whereas the hemodynamic parameters exhibited a significant time-of-day dependent effect. We conclude that these findings urge the need to account for the time of data acquisition in future MRI studies.

## Introduction

During the last two decades there has been an exponential increase in the number of publications related to brain functional connectivity (FC), as measured by functional magnetic resonance imaging (fMRI) (Pawela & Biswal, 2011). To date, research on FC has boomed covering a variety of disciplines, such as neurology, psychiatry, and oncology (Fox & Greicius, 2010; Woodward & Cascio, 2015; Bruno, Hosseini, & Kasler, 2012; Raichle, 2015). Resting-state functional connectivity (rs-FC) measures the temporal correlation of a spontaneous BOLD signal among the different brain regions. Even though the rs-FC networks have been found to be stable across population (Damoiseaux et al., 2006), the literature indicates vast individual differences based on a number of traits. Current evidence suggests that the changes or disruptions in the rs-FC may serve as a biomarker of brain disease, such as dementia (Broyd et al., 2009) and other neurodegenerative diseases (Brier et al., 2012). Deficits in cognitive performance associated with Alzheimer’s disease, mild cognitive impairment, autism spectrum disorders, and schizophrenia are also reflected in the rs-FC (Zhou et al., 2013, Hull et al., 2016, Sheffield & Barch, 2016). A recent publication on large scale UK Biobank data reported that differences in cognitive performance in healthy individuals are associated with differences in the rs-FC, specifically DMN, where the neural associations are also shared with individual differences in educational attainment and household income (Shen et al., 2018). Finally, the stimulus-related BOLD responses in most brain areas are found to change throughout the course of the day, with a typical decrease from morning to evening hours (Marek et al., 2010). The presence of diurnal brain dynamic is evident in literature concerning attention, which urges the notion that psychological and neuropsychological assessments together with work and school schedules, should instead be programmed in accordance with circadian rythmicity, age, and individual chronotype, rather than based on social and economic considerations, as the former are not easily adjusted (Valdez, 2019). Studies applying forced desynchrony protocol confirm that synchronized alternations between bursts of action potentials and periods of membrane hyperpolarization of cortical neurons are directly modulated by endogenous circadian rhythmicity (Lazar, Lazar, & Dijk, 2015). Consequently, chronotype markedly influences time-of-day modulation on cerebral activity patterns, where studies suggest that larks and owls exhibit an inverted relationship curve throughout the day (Christie & McBrearty, 1979; Horne, Brass, & Petitt, 1980). Notably, brain imaging studies using cognitive performance suggest that some but not all tasks vary throughout the day. For instance, insight-based problem performance is shown to increase at “non-optimal” time-of-day, contrary to the performance of solving analytical problems (Wieth & Zacks, 2011).

In spite of the endogenous nature of circadian rhythms in several brain functions, the time-of-day dependent variability in resting-state fMRI is not consistently reported. Functional connectivity in the DMN is reported to exhibit a rhythmic pattern, with its peak in the morning and lower in the later hours of the afternoon (Blautzik et al., 2013). Additionally, the variability in rs-FC has been reported in the medial temporal lobes (MTL) when comparing morning and evening scans (Shannon et al., 2013). When the magnitude of cerebral blood flow and functional connectivity in the DMN is examined, a consistent decrease in DMN functional connectivity, particularly in the posterior cingulate cortex (PCC) and the medial prefrontal cortex (mPFC) across the daytime is found (Hodkinson et al., 2014). Recent reports do support the notion that functional connectivity in the DMN is affected by time-of-day and chronotype (Facer-Childs, Campos, Middleton, Skene, & Bagshaw, 2019) and a steady decrease of global signal fluctuation and regional BOLD fluctuations together with whole-brain rs-FC throughout the day (Orban et al., 2020). Orban and colleagues (2020) further present evidence that the association between time-of-day and the reductions in global rs-FC is stronger than the association with fluid intelligence measure. Given the above, it is evident that the diurnal variation in rs-FC networks is present.

Further evidence for the circadian rhythmicity effect on neural measures is put forward by studies investigating cortical excitability, white matter microstructures, brain volume, grey matter density, cortical surface, and thickness (Voldsbekk et al., 2020, Karch et al., 2019, Nakamura, Brown, Narayanan, Collins, & Arnold, 2015; Huber et al., 2013; Trefler et al., 2016). Nakamura and colleagues (2015) report that brain volume changes significantly across the day, with larger brain volumes in the morning compared to the evening. These changes in brain volume are observed across multiple populations, i.e. healthy elderly individuals, patients suffering from multiple sclerosis, mild cognitive impairment, and Alzheimer’s disease (Nakamura et al., 2015). Morphometry measures suggest that increased volumes of cerebrospinal fluid CSF are associated with decreased volumes of grey and white matter (Trefler et al., 2016) and that the extracellular space volume is reduced in large parts of the white matter from morning to evening (Voldsbekk et al., 2020). It is suggested that the volume changes might also be associated with the level of individual hydration, where levels of hydration have previously been shown to have an effect on evoked BOLD signal captured by fMRI (Kempton et al., 2011). Given the existing body of literature, it is reasonable to question whether the captured individual differences in a healthy population, or group differences in clinical neuroimaging literature are indeed independent from the timing of the acquired scan (Trefler et al., 2016; Voldsbekk et al., 2020; Karsch et al.,2019). It is not recognized as a common practice to report the image acquisition time in neuroimaging studies, therefore it remains a possibly biasing factor. Exploring the effect of time-of-day in neuroimaging would contribute to a more robust reporting of data collection procedures, which would allow for an easier replication of findings if found significant.

More recently, a new approach to analyse fMRI data has been developed by Friston and colleagues (2010). Dynamic Causal Modelling (DCM), contrary to mainstream fMRI analysis techniques, incorporates Bayesian framework in order to estimate the non-linear relationship between brain regions, attempting to explain the connectivity based on predicted hidden neural states (Friston, Harrison, & Penny, 2003; Friston et al., 2014). Application of DCM in not limited to rs-FC, however it requires a predefined region of interest (ROI) and is data driven (Friston et al., 2003; Friston et al., 2014). Based on the rs-FC time-series, DCM generates measures of effective connectivity – i.e. a simulated directional relationship between the selected ROIs –– and separately models the hemodynamic parameters (Balloon model), amplitude (α), and spectral density (β) (Friston et al., 2014).

Given all the above, the aim of the current study was to investigate diurnal (time-of-day effect) change in resting-stated effective connectivity, measured by DCM. For the scope of this study, three large scale rs-FC networks were selected: the DMN, Saliency Network (SN), and Central Executive Network (CEN). The network effective connectivity was compared at six different timespans throughout the day (from 09:00 until 21:00). It was expected to observe diurnal change in neuronal activity and/or metabolic response (as generated by DCM).

## Methods

The Human Connectome Project data release “S1200” was used in the current study. For complete information about the dataset please see Van Essen et al., 2013 (http://www.humanconnectome.org).

### Participants

For the purposes of the current study, the participants scanned from 9:00 to 21:00 were selected from the complete dataset “S1200”. This decision was made given that only a few participants were scanned before 9:00 and after 21:00. The aim was to have an equal sample size for each timespan with equal gender distribution for each timespan. The mid-scan time was used to allocate the participants to 6 groups of selected timespans (9:00 h − 10:59 h, N=96; 11:00 h − 12:59 h, N=100; 13:00 h − 14:59 h, N=100; 15:00 h − 16:59 h, N=100; 17:00 h − 21:00 h, N=98), in total 594 participants (310 females).

All participants were healthy adults ranging from 22 to 35 years of age. The data was collected over 3 years on a single 3 Tesla (3T) scanner. The participant pool primarily consisted of subjects living in Missouri, in families with twins. Exclusion criteria were neurodegenerative disorders, documented neuropsychiatric disorders, neurological disorders, high blood pressure, and diabetes. For a complete list of inclusion criteria please see Van Essen et al., 2013 (Table S1)).

### Procedure

Participants were scanned at the Washington University (U.S.) over a two-day and one-night visit. An informed consent was signed by the participants at the beginning of day 1. In accordance with previously run pilot studies, a consistent scanning schedule was maintained for all the participants in the study, unless a re-scan was required. For more details, see Van Essen and colleagues (2013). A mock scanning trial with feedback on head motion was run prior to the first scanning. The data collection scanning for each participant was scheduled once a day with two rs-fMRI acquisitions each – one on day 1 (left/right and right/left phase encoding) and the other on day 2 (left/right and right/left phase encoding). The average time for each scanning occasion was 14.4 minutes, during which time the room was darkened. The participants were asked to lay still with their eyes open and fix their gaze on a bright fixation cross in the darkness. The clock time for each scanning day varied from 7:00 h to 22:00 h. Complete data collection procedure can be found in Van Essen et al., 2013. For the purposes of the current study, the averaged scan was used from either 1^st^ scanning session or the 2^nd^ scanning session based on the time of scanning acquisition.

### Image Acquisition

Resting-state fMRI data collection was carried out in accordance with an optimized fMRI image acquisition protocol as determined by the Human Connectome Project piloting. A custom Siemens 3T “Connectome Skyra” scanner was used to record the data for all participants. The scanner was equipped with a 32-channel head coil, custom gradient coils and gradient power amplifiers boosting the gradient strength to 100 mT/m. Resting-state fMRI data was acquired in two sessions: firstly 2×14.4 minute runs, Right/Left and Left/Right phase encoding on day one, and subsequently 2×14.4 minute runs Left/Right and Right/Left phase encoding on day two, a total of 1 hour resting-state fMRI. A gradient-echo multiband EPI imaging sequence was used to acquire resting-state fMRI data. Resting-state fMRI image acquisition settings were as follows: repetition time (TR) of 720 ms, echo time (TE) of 33.1 ms, 52° flip angle, field of view 208×180 mm (readout x phase encoding), slice thickness 2.0mm; 72 slices; 2.0 mm isotropic voxels and a multiband factor of 8 (for more information see Glasser et al., 2016; Uğurbil et al., 2013). In addition, high resolution T1-weighted structural images were obtained with the following parameters: TR 2400 ms, TE 2.14 ms, inversion time (TI) 1000 ms, flip angle 8°, FOV 224×224mm and 0.7 mm isotropic voxels (Uğurbil et al., 2013).

### Image processing

For the purpose of this study, the data was acquired pre-processed in accordance with “Human Connectome Project minimal pre-processing pipeline” (please refer to Glasser et al., 2013 for details). Standard pre-processing steps – such as correcting for distortions and spatial alignment – were performed. In addition to the mentioned standard procedures, the data was also corrected for spatial distortions, aligned across modalities and brought into a standard spatial atlas coordinate system. The only variation from standard pre-processing procedures was due to the bore diameter of the scanner being 56 cm, which is smaller than the standard Siemens 3T Skyra size (70cm diameter). The reduced diameter and lack of a customized patient table resulted in a higher placement of patient table in the bore and the participants’ heads not being centred along the gradient isocentre, meaning the scans have greater than normal gradient distortions, which have been corrected for in the Human Connectome Project pre-processed data used in this project, for more details see Van Essen et al., 2013.

### Analysis

To remove other nuisance confounds, the minimally pre-processed data were further processed prior to the DCM analysis. Using linear general models, as implemented in SPM12, the effects of the head movement (6 movement parameter) and the signal from white matter and cerebrospinal fluid areas were regressed out from the time series. This procedure has been performed separately for each of the two rs-fMRI sessions (Left/Right and Right/Left phase encoding, hereafter referred to as LR/RL) and for each individual separately.

There were 8 regions of interest selected for the current study: four nodes in the DMN (mPFC; PCC; right inferior parietal cortex, RIPC; left inferior parietal cortex, LIPC), two nodes of saliency network (SN) (anterior insula, AI; anterior cingulate cortex, ACC), and two nodes of central executive network (CEN) (dorsolateral prefrontal cortex, DLPFC; posterior prefrontal cortex, PPC). For exact coordinates of each ROI please see appendix 1. Each ROI was a sphere 6 mm in diameter. Cross-spectral-density dynamic causal modelling (csdDCM) implemented in SPM 12.2 was used to extract time series for each ROI (Friston et al., 2014). The 8 nodes were selected due to the computational power limitations.

The effective connectivity was estimated for each participant and for the LR and RL session separately. The PEB (Parametric Empirical Bayes) framework was used to estimate the joint effective connectivity per timespan (Zeidman et al., 2019).

A hierarchical PEB procedure was applied to test varying design-matrices against the time-of-day model. First, a PEB analysis was conducted for each timespan and for LR/RL separately. Thereafter, the corresponding twelve PEB results (6 timespans, 2 sessions) were subjected to a series of second level PEB analyses, where 18 models for time-of-day effects were explored. The models were conjointly specified for LR/RL, such that the two sessions were finally combined at this level of the analysis. The following models were specified:

1. Model 1-6: expectation of a *deviation from the mean for one single timespan*
2. Model 7-11: expectation of a *deviation from the mean over two adjacent timespans*,
3. Model 12-17: expectation of *phase-shifted variants of a sinusoid function* – approximating a circadian rhythm, peaking at different timespans
4. The null-model: expectation of *no predictions at any time span*

These 18 models were defined as different design matrices, comprising one column for the overall mean, one column for the mean-corrected model as described above, and five columns indicating the different timespans and for parametrising the repeated measurements, i.e., combining the LR and RL sessions. The results were compared using Bayesian model comparison (BMC). The winning model was explored at a cut-off of posterior probability of *P*p > 0.95. These analyses were conducted for the effective connectivity matrix (A-matrix, 8 x 8 parameter).

Then, the same procedure was repeated for the hemodynamic parameters of the Balloon model; *transit time*, *epsilon* and *decay* and the parameters α (reflecting amplitude) and β (reflecting the spectral density of the neural fluctuations). The parameter *transit time* was estimated for each of the eight ROIs, and *decay*, *epsilon,* α, and β are global parameters.

## Results

The BMC of the time-of-day variations in the effective rs-FC showed that the highest model evidence (model accuracy minus model complexity) was the Null model (see ****Figure 1a****). The effective connectivity matrix of the Null model with a posterior probability > 0.95 (pp; the updated probability of the model being true after comparison with other models) is displayed in ****Figure 1b****. In other words, the Bayesian model suggests that the neural activity in large scale networks remains stable throughout the day.

**Figure 1:**
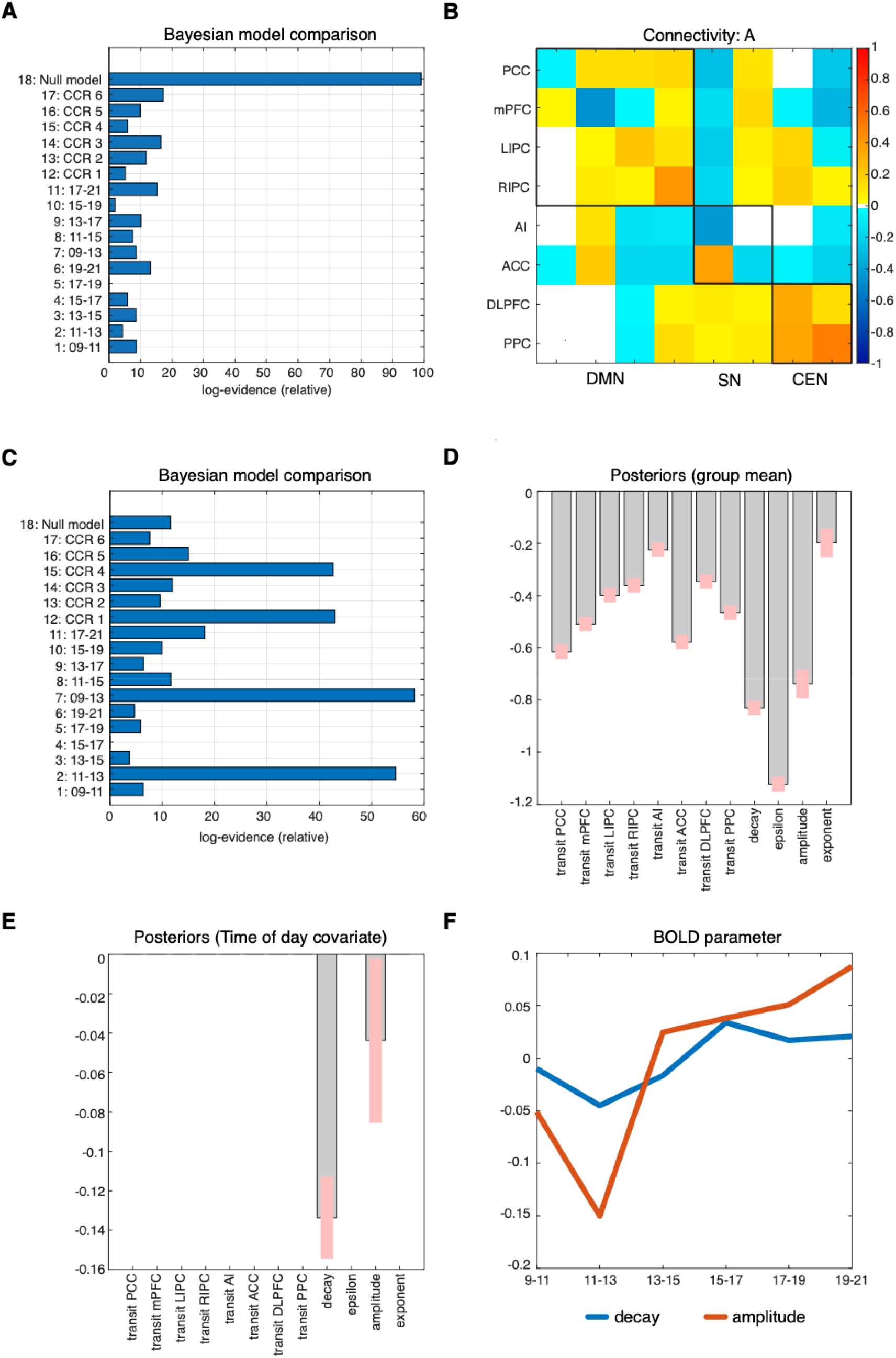
The figure summarizes all results from the hierarchical Parametric Empirical Bayes analysis. *a.* Bayesian Model Comparison on the connectivity parameter, where model 1-6 assumes deviating effect only for single time-spans of two hours, while model 7-11 assumed deviating effects for time-spans of four hours. The model 12-17 modelled different phase shifted version of an idealized circadian rhythm. Model 18 was the null model that assumed no time-of-day effects *b.* The estimation of effective connectivity (from columns to rows) across all subjects. The leading diagonal elements represent self-connections in logarithmic-scale relative to the prior mean of −0.5 Hz. White space represents no significant effect at pp level > 0.95 *c,* Bayesian Model Comparison on the hemodynamic parameter *transit time*, *decay, epsilon*, as well as *Cross-spectral-density amplitude* and *Cross-spectral-density exponent* *d.* Group means for the posterior estimates of the hemodynamic parameter, displayed at a posterior probability of pp>0.95 *e.* Posteriors of the winning model, displayed at pp>0.95 *f.* Time course of the two significant posteriors decay and CSD amplitude after time-span-wise averaging with Bayesian Model Averaging

Performing a model comparison on the hemodynamic parameters (*transit time*, *epsilon* and *decay*, as well as amplitude (α) and spectral density (β)), BMC favoured model 7, which predicted a deviation from the mean for the two timespans early in the morning, 09-13 h (see ****Figure 1c****). Moreover, three other models showed high model evidence, which were related variants of the winning model by predicting a deviation between 11-13 h (model 2) or they were a sinusoid function peaking between 11-13 h (model 12) or being the inverted wave form with a minimum between 11-13 h (model 15). Group means relative to the fitted model are shown in ****Figure 1d****. The posterior values seen in ****Figure 1e**** represent the level of parameter movement in accordance with the winning model (model 7) at the pp>0.95 level.

The posterior values for the *decay* and the *amplitude* of the CSD function are significantly reduced for the timespan 9-13 h (model 7). Finally, ****Figure 1f**** displays how the posterior values for decay and amplitude vary over the day, after the respective DCMs were averaged for each timespan using Bayesian Model Averaging – in other words, the neural metabolic response measured by BOLD. Using the parameters for the winning model, the effect sizes for *decay* and CSD *amplitude* are d=0.23 and d=0.28, respectively.

## Discussion

To the best of our knowledge, this study is the first to investigate the time-of-day influence on resting-state neuronal activity (effective connectivity) and metabolic demand (BOLD signal). The main findings of the current study are that while the functional- and effective connectivity of the brain remains stable throughout the day, the hemodynamic parameters defining the BOLD as generated by the DCM exhibit a variation between the selected timespans.

The effective connectivity in three large-scale resting-state networks, namely DMN, CN, and CEN, was found to remain stable across morning, noon and evening hours (from 09:00 to 21:00). These findings are contradictory to previously reported results that indicate a shift in DMN and MTL FC from morning to evening (Blautzik et al., 2013, Shannon et al., 2013). Interestingly, a recent publication using the same subjects (Human Connectome Project release S1200) reports a cumulative global signal (GS) decrease and whole brain FC decrease throughout the day (Orban et al., 2020), which appear as inconsistent with present findings of constant FC in DMN, SN and CEN. However, the conflicting findings between the current and previously reported results are likely rooted in different methodological approaches used that aimed to separate metabolic factors from connectivity. In the present study, the dynamic relationship between functionally connected nodes was assessed using a generative model, namely Dynamic Causal Modelling, which analyses the bidirectional effect each network exhibits with itself and others. By contrast, previous publications reported the functional connectivity, namely the correlational relationship between the nodes of the network, which is based on the temporal correlations of the BOLD signals (Facer-Childs et al., 2019; Orban et al., 2020). One might speculate that those correlative approaches are potentially biased by systematic variations of the underlying BOLD signal, as demonstrated by the present study. The main finding of our results is that the parameters defining BOLD signal, i.e., *decay* and *amplitude,* indeed show a time-of-day effect (model no. 7), and, hence, methods that are based on directly analysing the amplitude of the BOLD signal may be affected by the time-of-day effects. These effects might be caused by, for example, diurnal variations in the blood pressure (Millar-Craig, Bishop & Raftery, 1978).

The BOLD signal, according to the current findings, varies depending on the time-of-day, in particular between morning and afternoon, regardless of the effective network activity. The hemodynamic parameter *decay* and the *amplitude* of the CSD function all exhibited a significant relationship with the time-of-day dependent model (model no. 7) (****Figure 1e****). The *decay* parameter reflects the rate of signal elimination (Friston et al., 2000). A decrease in this parameter indicates an increased resting cerebral blood flow (rCBF) and a faster elimination of the signal and may also cause a larger BOLD undershoot. Accordingly, the BMC-selected model predicts a faster decaying BOLD signal before noon and, subsequently, a slower BOLD signal decay in the afternoon. As the plot over time indicates (****Figure 1f****), this appears to be a dynamic effect over the daytime, and the inclusion of further timespans may show a cyclic effect. A similar effect can be seen when examining the *amplitude* of the cross-spectral-density function that also demonstrates a lower amplitude in the morning than afternoon. In contrast to the *decay*, CSD *amplitude* showed a pronounced minimum for the 11-13 timespan. This different temporal evolvement during the morning does also explain why all four models that showed high model evidences in BMC (****Figure 1c****) were rated similarly. Interestingly, there were no region-specific effects in the *transit times*, which indicates that this is a global vascular effect. One might also speculate whether these globally appearing effects in *decay* and CSD *amplitude* might also explain the observed time-of-day dependent fluctuations of the global signal as observed by Orban and co-workers, using data from the Human Connectom Project, as well (Orban et al., 2020). It should be noted that the effect sizes are considerably large.

The shift of *decay* is in line with the studies on resting state, where the connectivity appears higher in the morning compared to the evening (Hodkinson et al., 2014). The time-of-day dependency of the BOLD signal as observed in varied *decay* and *CSD amplitude* parameters, is supported by previously published results on brain volume (Karsch et al., 2019, Nakamura et al., 2015; Trefler et al., 2016). The increase in *decay,* as seen in the current study occurs in the afternoon as well as the previously reported increase of CSF in Trefler and colleagues (2016). Given that CSF is vital for regulating the CBF, it is plausible to speculate that the reduction in the rCBF caused by a peak in the timespans before noon (9-11 until 11-13) is associated with the lower levels of CSF. During the first half of the daytime, a decrease in blood pressure, heart rate, and CBF(V) is observed around 11-13, whereas cortisol is still high. These daily changes in blood parameters provides support for the described change in the hemodynamic parameter. It is reasonable to observe a slower BOLD response with the reduced CBF, and it is likely that the *decay* is dependent on vascular signalling and relaxation time (Friston et al., 2000).

The authors of former publications controlled for chronotype, sleep duration, and quality (Blautzik et al., 2013; Shannon et al., 2013). In contrast, the current work was carried out on open access data that did not include information on chronotype or sleep depth. Even though the database includes the information regarding “bed time” and “get up time”, these are not considered to be accurate measures for the chronotype as they are driven by social and economic obligations – for example work/school hours – hence not measuring the indivudal free running sleep-wake cycles. A recently published work by Facer-Childs and colleagues (2019) suggests that there are significant differences in functional connectivity between early and late chronotypes in the DMN (Facer-Childs et al., 2019). These differences are somewhat in line with early findings of Merrow et al. (2005), and Weitzman et al. (1971), suggesting that different chronotypes have distinct rhythmicity patterns throughout the day, where the post-lunch dip and morning rise in cortisol levels might differ (Merrow et al., 2005; Weitzman et al., 1971). Taking into account these reported significant differences, it might be plausible that the reported effects from the current study could have been even stronger, had accounting for different phenotypical profiles been possible.

There are some limitations to the current work which could be addressed in future studies. First and foremost, the data obtained from the Human Connectome Project, despite being high in quality, was not collected in order to address the connectivity differences across the time-of-day and did not account for chronotype differences. Secondly, the analysis method used in the present study (DCM) differentiates between the parameters defining the hemodynamic response (BOLD) and connectivity measures, but no additional parameters that have been previously shown to affect resting state connectivity could be included, such as blood pressure, cerebral blood flow, respiration (Facer-Childs et al., 2019; Specht, 2019).

Furthermore, the study relied on between subject design, i.e., different subjects were randomly assigned to the timespans, whereas in order to obtain a truly comprehensive understanding of the circadian rhythmicity, a repeated measure (within subject design) would be best, where the same participant would undergo scanning at each timespan (Karch et al., 2019). Even though the individuals in the HCP were scanned twice (on day 1 and on day 2), the hours of image acquisition for the same participant differ between days, therefore limiting the present analysis to between subject design. Finally, in the present sample the family structure was not accounted for, which might have decreased the significance of lessened the results.

The fact that the significant time-of-day effect only occurred in the hemodynamic and not in the connectivity parameters has a critical implication for past, current, and future research studies. The results from research on group differences, which did not counterbalance or did not account for the time of scanning might have been in part detecting the time-of-day differences rather than the true group differences. For instance, when a group of participants is examined during morning hours (e.g. patients) and another group in the evening (e.g. control group). The use of dynamic causal modelling in future studies could aid in explaining the relationship between BOLD signal and FC. These future achievements would heavily contribute not only by bringing consensus in diurnal rhythmicity and resting-state connectivity research but also for in-depth understanding of brain activity.

In summary, the results from the current study contribute to the limited body of literature on time-of-day changes in the brain. The findings suggest that even though there is an observed variability during the course of the day in the hemodynamic response, which is captured by fMRI, the effective connectivity remains stable. The changes in the hemodynamic response possibly reflect the influence of time-of-day effects of the metabolic and vascular system. This may indicate that the BOLD signal is more susceptible to exogenous parameters than the brain activity itself. These findings urge the need for further separation between the hemodynamic response and neural activity as reflected by FC, since the relationship between the two might be more complex than previously thought and finally not stable throughout the day. Further, the time-of-day dependent variation of the metabolic basis of the fMRI signal might also partly explain the low reliability of fMRI studies, given that the effect size is rather large an effect that is typically not accounted for.

